# Conversion of iPS derived hepatic progenitors into scalable, functional and developmentally relevant human organoids using an inverted colloidal crystal poly (ethylene glycol) scaffold engineered from collagen-coated pores of defined size

**DOI:** 10.1101/296327

**Authors:** Soon Seng Ng, Kourosh Saeb-Parsy, Joe M Segal, Maria Paola Serra, Samuel J I Blackford, Marta Horcas Lopez, Da Yoon No, Curtis W Frank, Nam Joon Cho, Hiromitsu Nakauchi, Jeffrey S Glenn, S Tamir Rashid

## Abstract

Generation of human organoids from induced pluripotent stem cells (iPSCs) offers exciting possibilities for developmental biology, disease modelling and cell therapy. Significant advances towards those goals have been hampered by dependence on animal derived matrices (e.g. Matrigel), immortalized cell lines and resultant structures that are difficult to control or scale. To address these challenges, we aimed to develop a fully defined liver organoid platform using inverted colloid crystal (ICC) whose 3-dimensional mechanical properties could be engineered to recapitulate the extracellular niche sensed by hepatic progenitors during human development. iPSC derived hepatic progenitors (IH) formed organoids most optimally in ICC scaffolds constructed with 140 µm diameter pores coated with Collagen in a two-step process mimicking liver bud formation. The resultant organoids were closer to adult tissue, compared to 2D and 3D controls, with respect to morphology, gene expression, protein secretion, drug metabolism and viral infection and could integrate, vascularize and function following implantation into livers of immune-deficient mice. Preliminary interrogation of the underpinning mechanisms highlighted the importance of TGFβ and hedgehog signalling pathways. The combination of functional relevance with tuneable mechanical properties leads us to propose this bioengineered platform to be ideally suited for a range of future mechanistic and clinical organoid related applications.

## 1. Introduction

Induced pluripotent stem cells offer exciting new possibilities in developmental biology, disease modeling and transplantation. Comprehensive realization of that promise is likely to require consolidation with other emerging technologies such as 3D cell culture. Along those lines, hepatic progenitor cells derived from iPSCs [1] and primary tissue [2] were recently shown to form “Organoids” (self-organizing miniaturized 3D structures resembling liver tissue) following 3D culture initiated by Matrigel. Organoid generation using Matrigel has similarly been demonstrated across a broad range of tissue types including intestine, kidney and brain [3–5], underlining the importance of this novel approach across the stem cell field. Downstream applications are however limited by the use of Matrigel which is poorly characterized, highly variable and of mouse origin [6]. A bioengineered substitute is therefore essential and was recently reported for intestinal organoid generation [7]. A similar bottom up engineering approach for liver organoid production is urgently needed. We recently developed a novel hepatocyte culture system composed of a 3D-hexagonally arrayed inverted colloidal crystal (ICC) scaffold [8]. In comparison to other 3D culture systems, the ICC boasts several advantages. These include being made from an FDA approved material, polyethylene glycol (PEG), that can be functionalized with select ECM proteins or varied in mechanical stiffness, a highly uniform architecture which incorporates size-selectable pores to facilitate tissue interconnection across the whole module whilst retaining uniform nutrient penetration to all cells and finally, being transparent to provide an easy means of viewing cells and intra-cellular fluorescence over time. Exploiting these properties, we were able to facilitate induction of organoid structures with advanced liver function in primary human fetal liver cells (PHFL) cultured in ICC’s [9]. Lack of donor material, ethical constraints and biological heterogeneity however make further progress difficult with PHFLCs. We therefore turned to the recently established technology of human iPSC-derived hepatocytes [10][11], as a good biological approximation to PHFLCs but not limited by the same constraints cited above, to explore their potential for organoid generation.

## 2. Materials & methods

### Study design

The overall objective of this study was to test whether organoids bioengineered in ICC scaffolds exhibited a more physiologically relevant liver phenotype compared to conventional 2D or 3D (Matrigel) platforms. iPSC-derived hepatic progenitors (IH) differentiated from the same parental line were therefore matured in three different in vitro models (ICC, 3D Matrigel, 2D) for 14 days and compared. Measurement techniques to compare the three models were designed to characterize two major endpoints: morphology (organoid formation) and function (gene expression, protein expression, liver specific functions and in vivo behavior). In transplant experiments, littermate animals were randomly assigned to experimental groups and analysis of samples was performed blindly. Studies were repeated at least three times.

### Cells

All human tissues were collected with informed consent following ethical and institutional guidelines. Freshly isolated hepatocytes were obtained from Triangle Research Labs (TRL) and Human Developmental Biology Resource (HDBR) University College London (UCL). Fetal hepatocytes were obtained from 16-20 week old fetuses (HDBR), dissociated as previously described [12]. Three different human iPSC lines were used to generate hepatic progenitors (IH) for these experiments: [1], [10], (https://stemcells.nindsgenetics.org/)[11] the latter of which is considered to be of ‘clinical grade’ and thereby potentially suitable for future human therapy.

### ICC fabrication

Sacrificial crystal lattices were constructed using polystyrene beads with diameter of 40µm, 60µm, 100µm, or 140µm (Duke Scientific) and annealed under 140˚C for 1.5 hours. Lattices were then infiltrated with precursor solution of 50 %w/v polyethylene glycol-diacrylate (PEGDA, Alfa Aesar), 10 %w/v acrylate-PEG-N-hydroxysuccinimide (Laysan Bio Inc, AL) and 1 %w/v photoinitiator (Irgacure® 2959, BASF, Switzerland) in DI water. The precursor solution underwent free-radical induced polymerization under 75 W xenon ultraviolet (UV) light source (Oriel Instruments, Mountain View, CA) for 10 min. After polymerization, the polystyrene sacrificial lattice was removed by tetrahydrofuran treatment for two hours. The resulting ICC hydrogel scaffolds were equilibrated in deionized water and conjugated with appropriate ECM proteins (Fig. 1A).

**Fig. 1.**
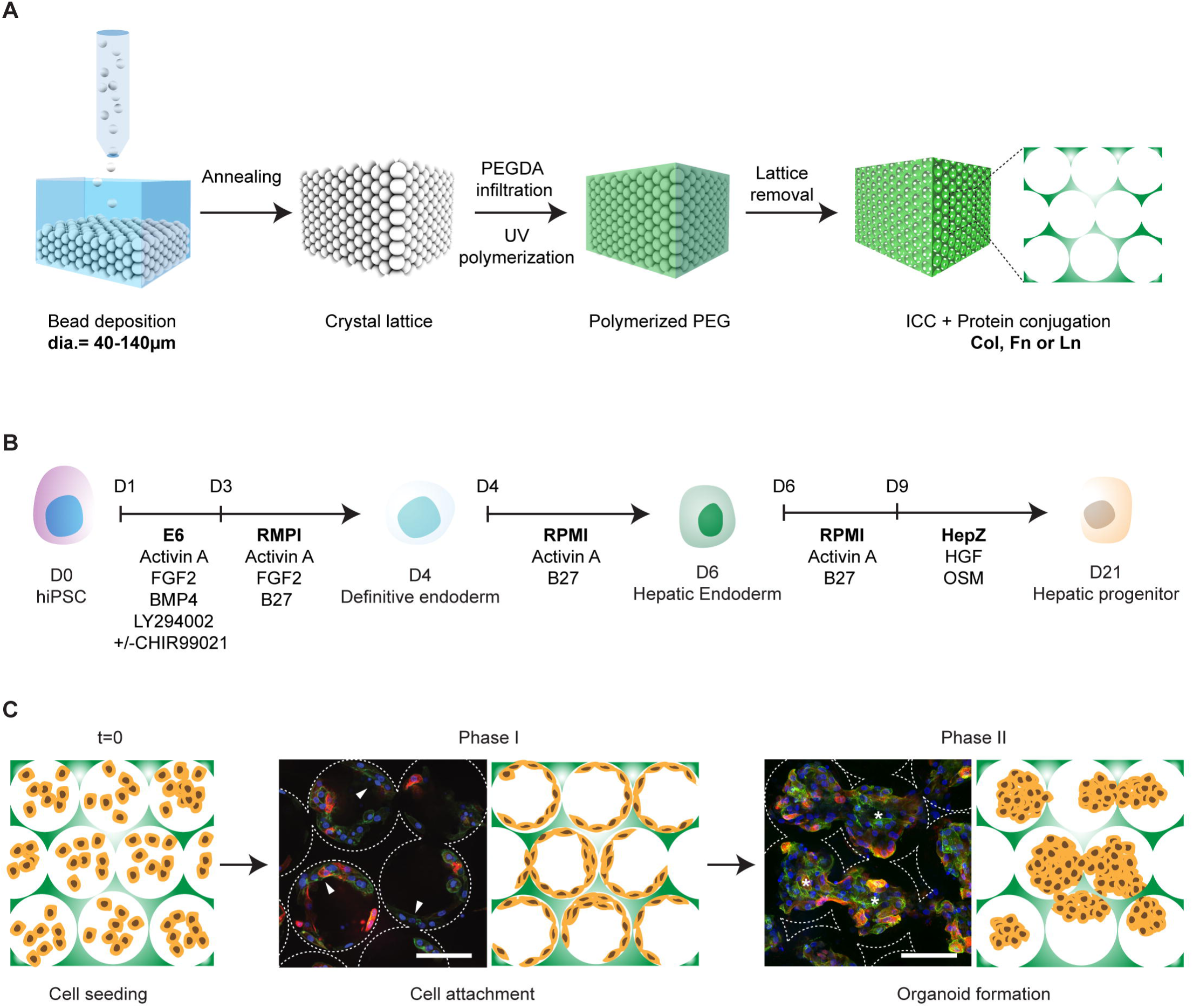
Bioengineering liver organoids using ICC scaffold and iPSC-derived hepatic progenitor (IH). (**A**) Schematic illustration of ICC fabrication using a range of sacrificial monodispersed beads and ECM proteins for stem cell niche modulation. (**B**) Schematic of human iPSC-derived hepatic progenitors differentiation protocol. (**C**) Schematic of bioengineering liver organoids using IH and ICC in three main steps, cell seeding, cell attachment (Phase I) and organoid formation (Phase II). Confocal micrographs of human fetal liver cells (Fetal; CTNNB green; CK19 red) demonstrating the two-phase organoid formation in ICC. Arrowheads indicate cells lining surface of ICC; asterisks represent cells forming clusters. Scale bar, 100µm.

### iPSC-derived liver organoid culture

iPSC-derived hepatic progenitors (IH) were generated using our established protocol (Fig. 1B) [13]. To generate iPSC liver organoids, we suspended approximately 0.5×10^6^ IH cells in Hepatozyme medium (Life Technologies), supplemented with Oncostatin M 0.01 µg/ml (Peprotech) and Hepatocyte Growth Factor 0.05 µg/ml (Peprotech) with the final cell density of 125×10^6^ cells/ml (Fig. 1C). Approximately 4µl cell suspension was then pipetted onto the surface of partially dehydrated ICC scaffolds and placed in an incubator for 30 min without media to minimize cell loss and maximize cell attachment. Cell-laden ICC scaffolds were then transferred to a new 48-well plate for culture. Cells were left for a further 14 days to mature. Media was refreshed every two days. 2D controls were generated using IH cells seeded into standard 48 well plate tissue culture plastic plates coated with the same ECM proteins and cultured with the same reagents as used for ICC culture.

### Immunofluorescence staining and imaging

After fixation with 4% paraformaldehyde, cells were blocked and permeabilized in 1% w/v bovine serum albumin (BSA, Sigma-Aldrich), 10% donkey serum (Life Technologies) and 0.1% Triton) for 30 min at RT. For nuclear antigens, cells were treated with 0.5% Triton (Sigma-Aldrich), then primary antibodies, CTNNB (mouse, 1:100; bdbiosciences), CK19 (rabbit, 1:500; abcam), HNF4a (rabbit, 1:100; abcam), AFP (mouse, 1:100; abcam) and ALB (goat, 1:100; Bethyl), ASGR1 (mouse, 1:100; Thermo Fisher Scientific), COL1 (rabbit, 1:500; abcam), MRP2 (mouse, 1:200; abcam), ZO-1 (mouse, 1:100; abcam) and CD26 (rabbit, 1:100; abcam) were applied for overnight at 4˚C. After washes, cells were then incubated with Alexa 647, Alexa 568, Alexa 488 conjugated secondary antibodies (Life Technologies). Samples were counterstained with DAPI (NucBlue, Life Technologies). Confocal micrographs were captured using Nikon Ti spinning disk confocal microscope equipped with Andor Neo camera and images were processed by NIS-element software.

### Cell number quantification

Double stranded DNA was collected, quantified and interpolated from standard curve plotted using a range of known cell number as illustrated by Quant-iT^TM^ PicoGreen^TM^ dsDNA Assay Kit (ThermoFisher Scientific)

### Human albumin Enzyme-linked Immunosorbent Assay (ELISA)

Albumin secretion of all different cell types was assessed using the Human Albumin Quantitation Set (Bethyl Laboratories Inc) following manufacturer’s instructions.

### Bright field imaging

Bright field imaging was captured using Leica DMIL LED equipped with Leica DFC3000 G camera and images processed by LAS X software.

### Immuno-histochemistry staining and imaging

IH-ICC or animal explants were fixed in 10% formalin buffer saline for two days then dehydrated and paraffin wax infiltrated using Excelsior™ AS Tissue Processor. After embedded, sectioned (5µm), and stained using Mouse and Rabbit Specific HRP/AEC (ABC) Detection IHC Kit (abcam) using antibody CK19 (rabbit, 1:200; abcam), EPCAM (mouse, 1:200; abcam) and ALB (mouse, 1:100; abcam) then counterstained with Eosin Y dye (abcam). Mounted slides were imaged using NanoZoomer (Hamamatsu).

### Fluorescence-activated cell sorting (FACS) staining and analysis

Cells were dissociated from organoid or 2D culture using TrypLE reagent (Thermo Fisher Scientific) and stained using the same condition as immunofluorescence staining protocol stated above. Flow analysis performed using FACSCanto II (BD) and analyized using FlowJo (LLC).

### Quantitative real-time PCR (RT-qPCR) analysis

Total RNA was harvested using TRIZOL reagent (Sigma), treated with DNase (Promega) and phenol/chloroform purified. For each sample 0.5 µg of total RNA was reverse transcribed using SuperScript VILO cDNA Synthesis kit (Thermo Fisher Scientific). A typical RT-PCR reaction contained 10ng of sample cDNA, 0.0075µl of individual forward and reverse primer each at 100µM stock, 5µl Taqman Universal Master mix (Applied Biosystems), 1ul Taqman target probe (supplementary Fig. 5) and made up to 10µl with nuclease-free water. Real time PCR reactions were amplified for 40 cycles on a CFX384 Touch™ Real-Time PCR Detection System (Biorad) in triplicate and normalized to ACTB in the same run.

### Heat map generation, gene set enrichment analysis, functional network analysis and top canonical pathway analysis

Heat maps were generated from Bulk RNAseq data collected from three Human Fetal Livers, three Human Adult Livers and three iPSC-derived hepatocytes harvested at definitive endodermal stage from the BOB cell line. The three fetal livers included 14pcw, 16pcw and 20pcw. The three adult Livers were Female 18yrs, Male 43yrs and Male 13yrs. RNA was extracted using TRI reagent. Starting with 1ug input total RNA, Ribosomal RNA was removed using Ribo-Zero Gold rRNA Removal kit (Illumia). Sequencing libraries were prepared using NEBNext® Ultra^™^ Directional RNA Library Prep Kit for Illumina (N.E.B) using 100ng rRNA depleted sample and sequenced on a HiSeq 2500 system in Rapid run mode (Illumina) following a standard protocol. All libraries generated between 15 and 25M reads. Reads were mapped to GRCh38 reference genome using Bowtie2. Raw counts and normalized gene expression was generated using HT-Seq and DESEq2 packages respectively. Heat map was generated using R (http://www.R-project.org) (R Development Core Team, 2008). Heat maps represent average DESEq2 normalized gene expression values of three independent biological samples. Gene set enrichment analysis was performed on normalized RNA sequencing gene expression data through GSEA software^27,28^ run using the hallmark MSigDB gene set collections. To examine the role of IH-ICC organoid in cell polarity and bile acid synthesis and metabolism, we cross-referenced the differential gene expression with the Gene Ontology (GO) annotations and predict the protein interactions networking using Search Tool for the Retrieval of Interacting Genes (STRING). Top canonical pathways analysis was performed using Ingenuity Pathway Analysis (IPA) (Qiagen).

### GLuc HCVcc infection and analysis

GLuc HCVcc stocks were prepared using electroporation on Huh 7.5 cells as described GLuc HCVcc is a full length Gaussia luciferase reporter virus Jc1FLAG2 (p7-nsGluc2A)35 infectious Jc1 genome that allows us to monitor viral infection in real time. Knock down HCVcc is the corresponding polymerase-defective mutant that inhibits viral replication. Viral stocks were prepared using electroporation on Huh 7.5 cells as described[14]. Infectious titers of HCVcc inocula were determined using Huh 7.5 cells as described. Cultures were inoculated for 3 hours with HCVcc then washed extensively with PBS and fed with culture media.

### Animal experiments

All animal experiments were performed in accordance with UK Home Office regulations (UK Home Office Project License number PPL 70/8702). Immunodeficient NOD.Cg-Prkdc^scid^ Il2rg^tm1Wjl^/SzJ (NSG) mice which lack B, T and NK lymphocytes^18^ were bred in-house with food and water available ad libitum pre- and post-procedures. A mix of male and female animals were used, aged approximately 6-8 weeks. An incision was made in the capsule of the caudate lobe of the liver and a ‘pocket’ was created by raising a flap of capsule on either side of the incision. The scaffold was slid into the pocket and covered by the capsular flaps. The left lobe of the liver was then allowed to cover the site of the transplant.

### Statistical Analysis

N in the paper represents the number of biological replicates of each batch of liver specific differentiation performed using three iPSC lines. Statical significance (p < 0.05) was determined by using Student’s T-test (assume Gaussian distribution, two-tailed) or One-way ANOVA followed by Tukey’s posthoc test.

## 3. Results

### 3.1. iPSC-derived hepatic progenitors (IH) form organoids analogous to fetal liver derived primary cells following culture in ICC scaffolds

Human fetal liver cells seeded into ICC scaffolds attach as a single cell layer (Phase I – days 0 to 3 post-seeding) before organising into morphologically stable, interconnected clusters (Phase II - days 7 post-seeding onward) (Fig. 1C). The resultant 3D structure resembles a mini-liver or hepatic ‘organoid’ with demonstrated advanced liver function [9]. We therefore hypothesized that organoid formation was critically dependent upon the translation of specific ‘cell-matrix’ and ‘cell-cell’ signals by hepatic progenitors in a manner which could be recapitulated using hiPSC-derived progenitors (IH) (**Supplementary Fig. 1**). To test this hypothesis, we therefore engineered a library of ICC scaffolds to evaluate the effects of selected extracellular matrix protein (ECM) coatings and pore sizes on IH-organoid forming ability (Fig. 2). Of the ECM coatings tested, only cells seeded into collagen (Col) were able to attach and self-organize into interconnected clusters as observed with primary fetal progenitors (Fig. 2A-C). Cells seeded into fibronectin (Fn) attached but failed to establish confluency whilst cells seeded into laminin (Ln) attached but failed to organize into interconnected clusters. The observed organoid forming ability was matched by functional parameters of liver tissue such as albumin secretion (Fig. 2D). Having established the optimal ECM coating, we next went on to assess the effects of cell-cell interaction by testing Collagen coated ICC’s composed from different pore sizes. Accordingly, a 140µm pore size was found to be optimal for organoid formation based on morphology and again confirmed by hepatic function (Fig. 1E-H and Supplementary Fig. 2). Through this extensive series of experiments, we observed IH seeded into Collagen coated ICC scaffolds of 140 µm pore size best replicated the results seen with primary fetal liver cells. Importantly, the two-phase process of organogenesis was observed only with IH and primary fetal cells but did not occur with primary adult or cancer-like hepatocytes (Huh 7.5 and HepG2) (**Supplementary Fig. 3**).

**Fig. 2.**
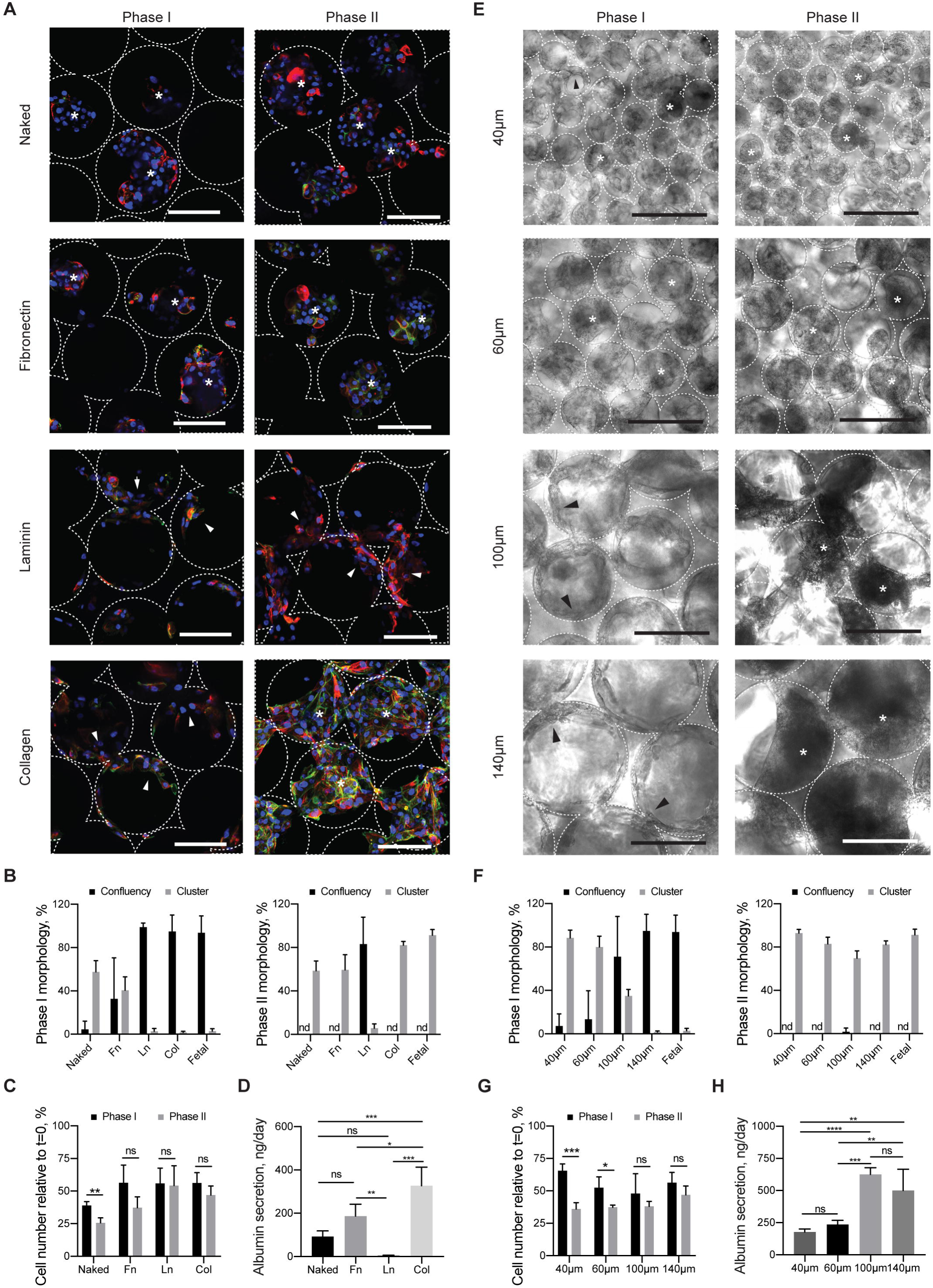
Characterizing the effects of pore size and ECM proteins on liver organoid formation in ICC scaffold. (**A**) Confocal micrographs of IH (CTNNB, green; CK18, red) following seeding into non-coated (Naked), Fibronectin (Fn), Laminin (Ln), and Collagen (Col) coated ICC’s. (**B**) Quantification of cellular surface lining (confluency) and cluster formation ability seen across different ECM’s during Phase I and II. (**C**) Cell number relative to initial cell seeding number in Phase I and II. (**D**) Albumin secretion rate of IH seeded in different ECM coated ICC in Phase II (**E**) Bright field images revealing the morphologies of IH in ICC with different collagen coated pore sizes (40µm, 60µm, 100µm and 140µm) and the respective (**F**) morphological quantitation and (**G**) relative cell number in Phase I and II. (**H**) Albumin secretion rate of IH seeded in ICC with different pore sizes in Phase II. Arrowheads indicate cells lining surface of ICC; asterisks represent cells forming clusters. Scale bar, 100µm. Mean±sd, N=4. ^*^p< 0.05; ^**^p< 0.005; ^***^p< 0.0005; ^****^p<0.0001; ns non-significant.

### 3.2. IH-ICC derived organoids demonstrate morphological and transcriptomic characteristics of human liver

To evaluate the anatomical similarity between organoids derived from iPSC-hepatic progenitors cultured in ICC scaffolds (IH-ICC) and human liver, we first used a combination of histochemical and immunofluorescence staining to characterise the organoid tissue across the two phases of organogenesis previously described (Fig. 3A-B and Supplementary Fig. 4-6). In Phase I, hepatic progenitor markers such as keratin 19 (CK19), epithelial cell adhesion molecule (EPCAM) and alpha-fetoprotein (AFP) proteins were expressed in cells lining the circumference of the scaffold’s pores. Following onset of organogenesis in Phase II, these (CK19, EPCAM, AFP) +ve progenitor cells were retained at the periphery of the organoid, with more mature cells, characterised by protein expression of asialoglycoprotein receptor 1 (ASGR1), albumin (ALB), hepatocyte nuclear factor 4-alpha (HNF4a) forming and remodelled by the cell secreted type I collagen (Col-I) at the centre. These data were then validated quantitatively using flow cytometry (Fig. 3C) and by RT-PCR for gene expression (Fig. 3D), allowing us to conclude that organoids became established with graded anatomical distribution of cellular maturity in a structure reminiscent of the hepatic buds seen in embryonic [15] and regenerating human liver tissue [16]. This anatomical distribution was not observed when IH were cultured in Matrigel, the current gold standard in the field. Embedding in Matrigel, IH established lumen-containing colonies, expressing mature hepatic markers (**Supplementary Fig. 7**) with functional properties (protein secretion and metabolic activity) significantly higher compared to 2D culture but notably not significantly higher than in ICC culture (**Supplementary Fig. 8**).

**Fig. 3.**
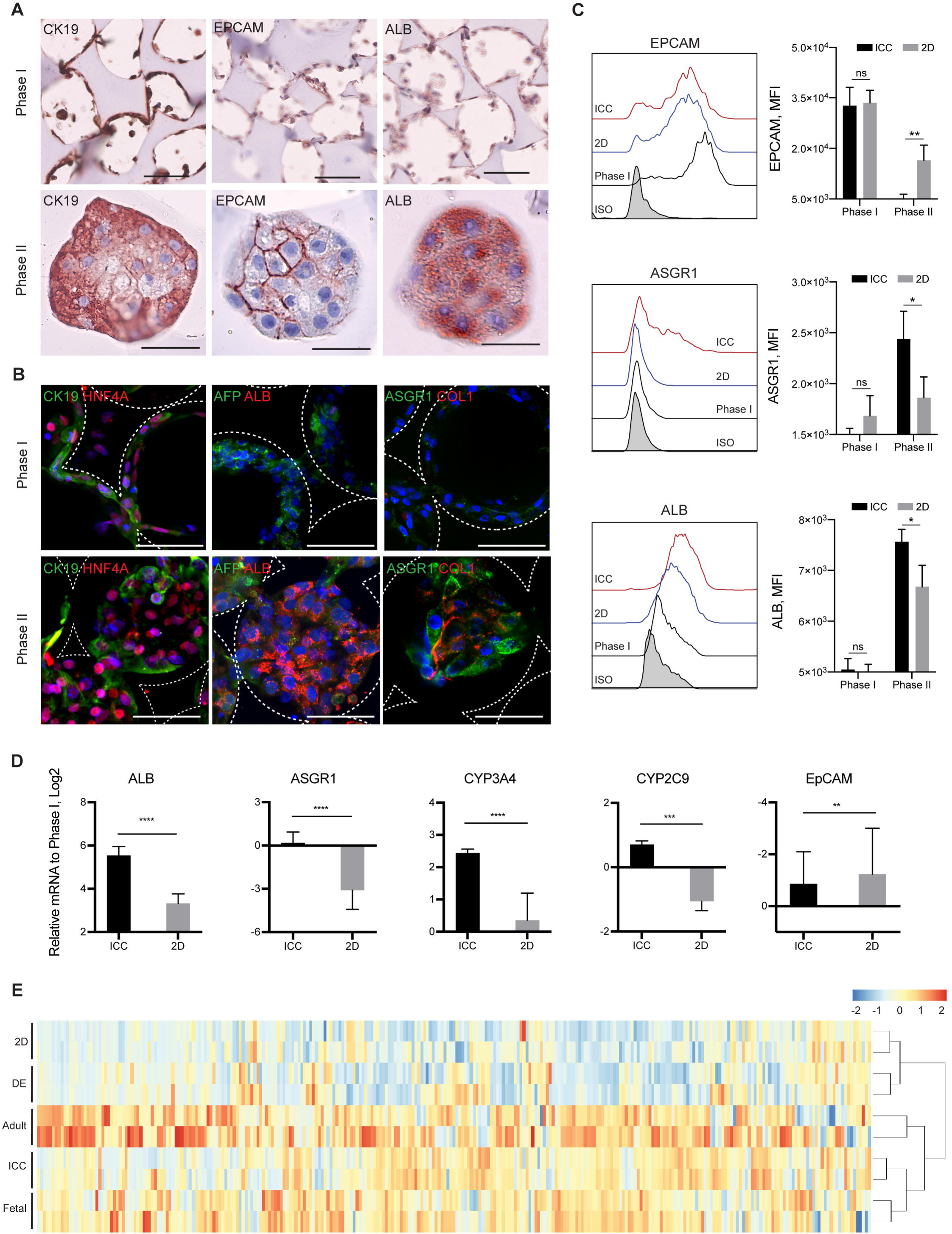
Morphological and transcriptomic characterization of IH-ICC organoids. (**A**) Histochemical images demonstrating morphogenesis of IH from a single cell layer (Phase I top panel; scale bar, 100µm) to organoids (Phase II bottom panel; scale bar, 50µm) occurs in conjunction with differential protein expression of developmental markers CK19 (left), EPCAM (middle) and ALB (right). (**B**) Confocal micrographs highlighting upregulated protein expression of mature (ASGPR1, COL1, ALB) hepatic markers occurs in conjunction with down regulation of immature (CK19 and AFP) markers during transition of IH from Phase I (top panel; scale bar, 100µm) to Phase II (bottom panel; scale bar, 100µm) organoids. (**C**) FACS histogram and mean fluorescence intensity (MFI) analysis demonstrating hepatic maturation kinetics of IH in ICC vs 2D culture (N=4). (**D**) Differential gene expression (by RT-PCR) of selected genes reveals a more mature hepatic signature of IH in ICC vs. 2D culture (N=8). (**e**) Bi-clustering heatmap of 296 liver-specific genes across different primary (adult & fetal liver) and IH (DE, 2D & ICC) samples. Samples are linked by the dendrogram above to show the similarity of their gene expression patterns. Mean±sd, *p<0.05; **p< 0.005; ***p< 0.0005; ****p< 0.0001; ns nonsignificant.

**Fig. 4.**
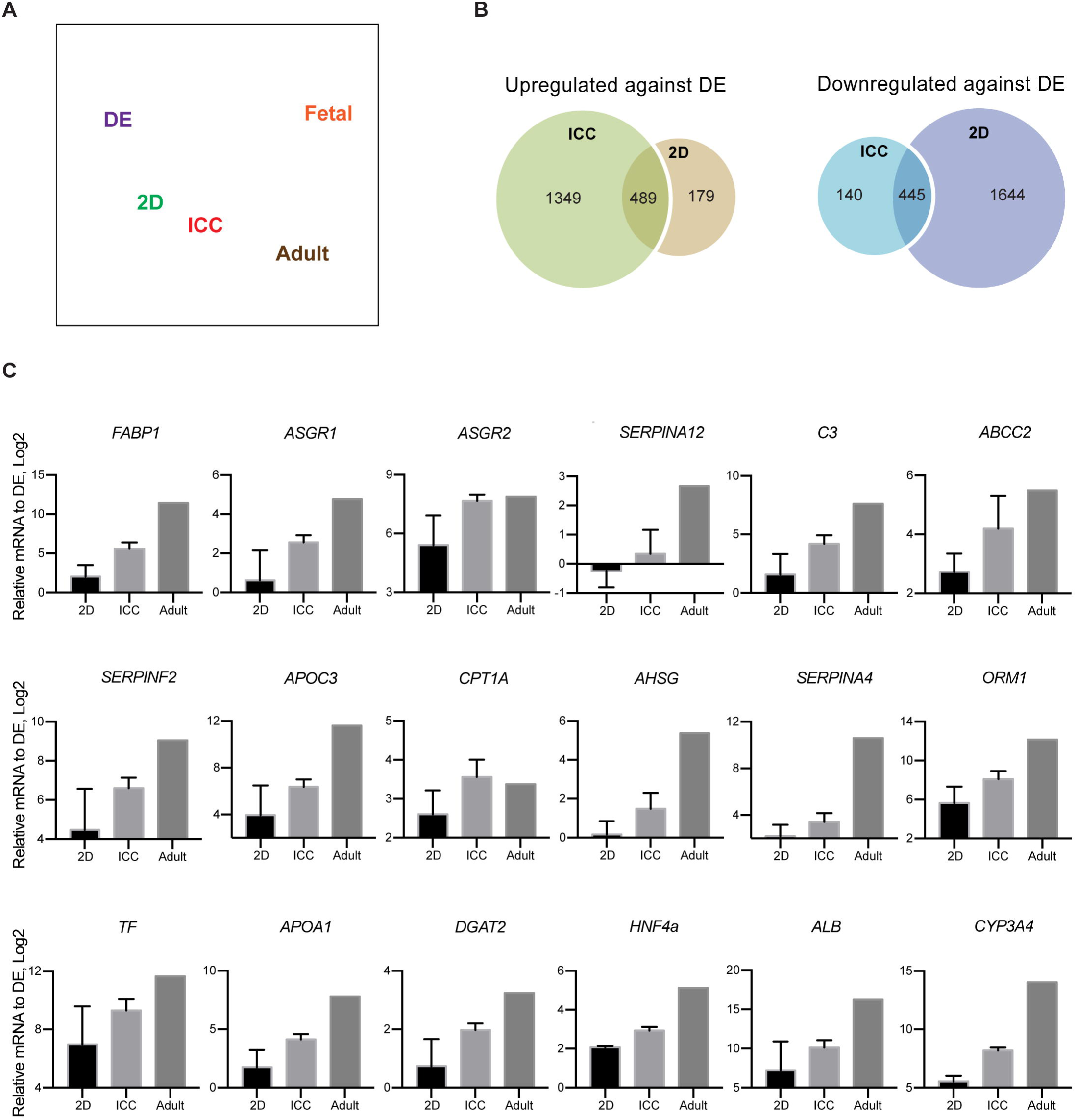
The transcriptomic analysis of liver organoid in ICC. (**A**) Principle component analysis of RNA-seq data demonstrating the proximity of gene expression variance in 2D plot. (**B**) Venn diagrams showing the number of up and downregulated gene in ICC and 2D with respect to DE. (**C**) RT-PCR validation on top 18 liver-specific genes identified by RNA-seq analysis (N=3) Mean±sd.

Having validated the organoid structure in our new bioengineered platform, we next sought to characterize the global transcriptomic changes during the different phases of organogenesis using RNA-seq. To do this, organoids were compared to iPSC derived definitive endoderm (DE), 2D cultured IH, primary adult and primary fetal hepatocytes. Principle component analysis (PCA) on DE, 2D, ICC, fetal and adult samples (Fig. 4A) shows that IH have a distinct transcriptional phenotype when cultured in ICC over 2D conditions. A liver-specific gene-set of 296 genes were selected to perform unbiased hierarchical clustering and visualized by heatmap. The unbiased hierarchical clustering revealed that the liver-specific transcriptional signature of organoids was positioned in the cluster next to adult and fetal liver, with higher similarity to fetal than adult, whereas 2D cells clustered next to DE (Fig. 3E). Overall 489 genes were found to be significantly upregulated in both organoid and 2D samples, with 1349 and 179 genes uniquely upregulated in organoid and 2D systems respectively (Fig. 4B). Gene set enrichment and Ingenuity Pathway (IPA) analysis of the upregulated genes revealed some liver-specific functions such as xenobiotic and bile acid metabolism to be enriched in both organoid and 2D platforms (**Supplementary Fig. 9 and Supplementary Table 1 and 2**) but the majority of liver-specific functions such as cholesterol homeostasis, glycolysis, fatty acid metabolism and protein secretion were found to be uniquely enriched in IH-ICC. The top 18 upregulated organoid specific hepatic genes were validated by qPCR (**Supplementary Table 3 and Fig. 4C**).

Cumulatively, these data confirm our previous morphological observations suggesting IH derived progenitors seeded into ICC scaffolds form organoid structures whose transcriptomic and protein expression profiles resemble human liver structures.

### 3.3. IH-ICC organoids are functionally closer to human liver

To interrogate their functional capability, we first evaluated albumin production rates of organoids over an extended period of time (Fig. 5A). We concluded that the increasing albumin production rate was principally a consequence of hepatic maturation rather than increase in cell number since a relatively quiescent proliferation rate was observed throughout the time window (Fig. 2C and G). Advanced hepatic function was then confirmed by increased basal activities of drug metabolizing cytochrome P450 isoforms CYP3A4 and CYP2C9 (Fig. 5B). Similarly, capability for transporting bile salts also appeared more advanced. Hepatic proteins critical for the structure of bile canaliculi such as multidrug resistance protein 2 (MRP2), tight junction protein 1 (ZO-1) and CD26 were all expressed (Fig. 5C and **supplementary Fig. 11**). Bile salt efflux was then validated through uptake and accumulation of synthetic bile acid, cholyl-lysyl-flourescence (CLF), in bile canaliculi-like regions (Fig. 5D). The ability of organoids to retain CLF was eliminated when treated with the BSEP inhibitor, Troglitazone (TGZ), as indicated by fluorescence measurement (Fig. 5E).

**Fig. 5.**
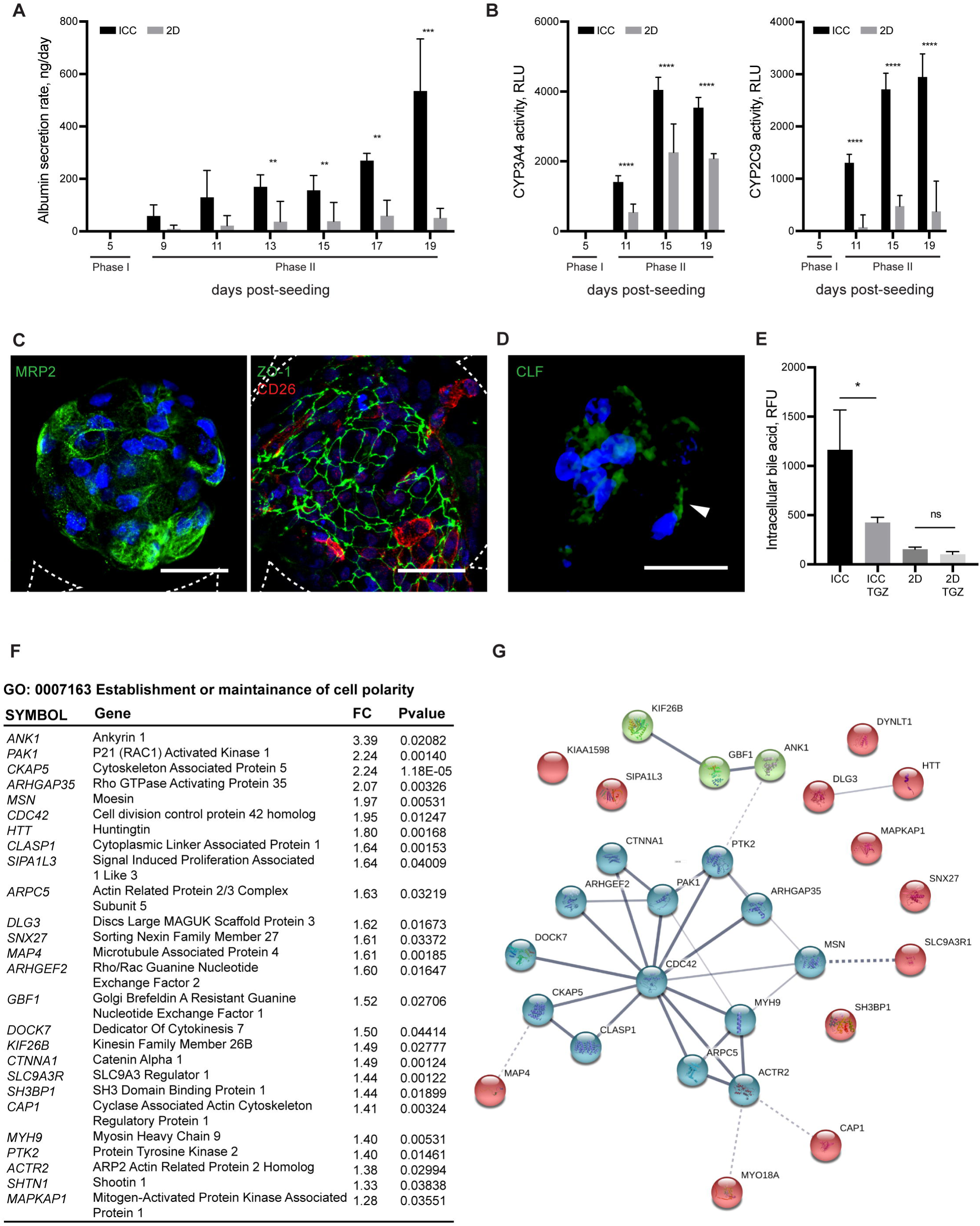
Functional validation of organoids. (**A**) Albumin secretion rate of IH-ICC vs. 2D (N=8). (**B**) Basal metabolic activity of Cytochrome P450 isoforms CYP3A4 and CYP2C9 in IH-ICC vs 2D **(N=8**). RLU, relative luminescence unit. (**C**) Confocal micrographs showing expression of hepatocyte polarity markers (MRP2, ZO-1 and CD26) in IH-ICC organoids. Scale bar, 50µm. (**D**) Accumulation of Cholyl-L-lysyl-fluorescein (CLS) in IH-ICC organoids after 40 minutes of CLF incubation followed by 40 minutes of washing. White arrowhead points to the CLF accumulation. (**E**) Effect of adding Troglitazone (TGZ) to CLS retention in IH-ICC (N=4). (**F**) A list of uniquely upregulated genes in IH-ICC vs 2D that involved in establishment and maintainance of cell polarity. FC, fold change. (**G**) The STRING functional network predicted the associations between proteins (nodes) from regulated genes involved in cell polarity in IH-ICC. The cluster analysis was performed using KMEANS clustering algorithms. Mean±sd, ^*^p<0.05; ^**^p< 0.005; ^***^p< 0.0005; ^****^p< 0.0001; ns nonsignificant.

The cell polarity and bile acid secretory function of organoid was further validated by cross-referencing the uniquely upregulated differentially expressed genes in ICC with GO categories (GO 0007163: establishment or maintenance of cell polarity; GO 0000902: cell morphogenesis; GO 0007010: cytoskeleton organization; GO 0006699: bile acid biosynthetic process and GO 0008206: bile acid metabolic process) followed by functional network analysis using STRING on selected enriched gene sets (Fig. 4F **and Supplementary Fig. 11-13**) [17]. The 28 genes uniquely upregulated in organoids could be assigned to the key KEGG pathways such as regulation of actin cytoskeleton, focal adhesion and tight junction required for cell polarization (Fig. 5G).

### 3.4. IH-ICC organoids are suitable for disease modelling and form vascularised tissue following transplantation

Next, to test the ICC organoid’s suitability for disease modeling we investigated whether organoids expressed HCV host factors and were susceptible to HCV infection. Transcriptional analysis revealed expression of HCV host factors were enriched in organoids and more closely resembled the expression signature of primary hepatocytes (Fig. 6A). This included genes responsible for HCV entry such as claudin 1 (*CLDN1*) occludin (*OCLN*) scavenger receptor, class B, type 1 (*SCARB1*), *CD81*, low-density lipoprotein receptor (*LDLR*) and heparin sulfate glycoprotein (*HSGP*). In addition, a wide range of apoliproteins responsible for packaging and interacting with viral particles were highly expressed [18]. Immunofluorescence staining showed CLDN1 appear to localize at the interface of cell-cell junctions, similar to ZO-1 and MRP2 (Fig. 6B **and Supplementary Fig. 14**). These results are in agreement with the CLDN1 distribution seen in HepG2 and Huh-7 cell models permissive to HCV entry [19, 20]. We next infected organoids with genotype 2a HCV reporter virus expressing secreted *Gaussia* luciferase (HCVcc) and knock-down HCVcc (kd-HCVcc) which is incapable of replication and acts as a negative control. Luciferase signal was only detected in organoids inoculated with HCVcc cultures, whilst 2D cells and kd-HCVcc inoculated samples failed to exhibit detectable signal (Fig. 6C).

**Fig. 6.**
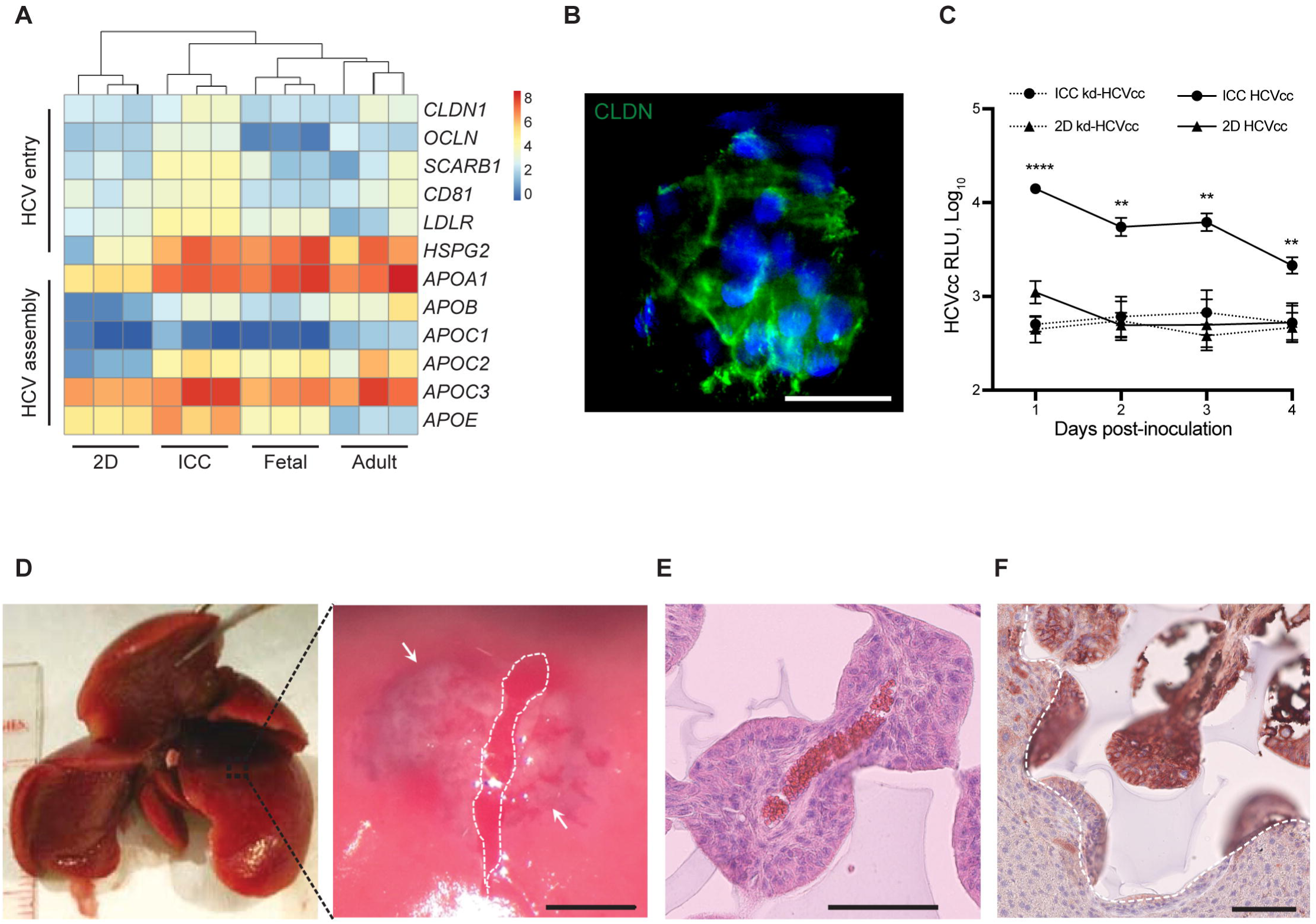
Disease modelling and in vivo transplantation. (**A**) Heatmap and hierarchal clustering comparing expression of 12 genes involved in encoding HCV entry and assembly in IH-ICC vs 2D vs primary (adult, fetal) liver. (**B**) Confocal imaging showing expression of claudin-1 (CLDN1) in IH-ICC organoids. Scale bar, 100µm. (**C**) HCV expression of IH-ICC vs 2D following infection with HCV reporter virus expressing secreted GLuc (HCVcc, N=4) or mock infected with knock down HCVcc (kd-HCVcc, N=3) and subsequently were sampled and washed daily. RLU, relative luminescence unit. (**D**) Photograph showing location of surgical pocket formation on murine caudate lobe (left) and appearance following IH-ICC transplantation (right). The white dashed line depicts the capsular incision and the limits of the sub-capsular scaffold implant are shown by the white arrows. Scale bar 1.5mm. (**E**) H&E staining of explant reveals neo-vasculature of IH-ICC. Scale bar, 100µm. (**F**) Immuno-histochemical staining of explant for human albumin. Dashed white line indicates the boundary between implant and host liver. Scale bar, 100µm. Mean±sd; ^**^p<0.005, ^****^p< 0.0001, nd not detected.

Having confirmed the organoid’s preferential suitability for drug metabolism and disease modelling we next sought to explore the effects of in vivo transplantation. A pocket on the caudate lobe of murine liver was created by making an incision in the liver capsule. Organoids were placed into this pocket and sandwiched in place between the left lobe and the lower caudate lobe in order to achieve a bona fide homeostatic environment (Fig. 6D). After 28 days, grafts were retrieved for further analysis. H&E staining revealed implants were well integrated into the host parenchyma, without evidence of significant fibrosis / inflammation whilst neo vascularization had successfully occurred between host and donor tissues (Fig. 6E). Histochemical staining with human albumin confirmed the implanted structures were of human origin, the organoid structure had remained intact and the presence of human albumin in host serum suggested cells remained functional (Fig. 6F).

### 3.5. TGFβ and Hedgehog Signalling Pathways are important for organoid formation

To identify signalling pathways involved in the orchestration of hepatic organoid formation, gene set enrichment analysis was performed as described before. The top 15 gene sets uniquely enriched in the ICC were related to metabolic/ biosynthetic and inflammatory/ immune related processes (Fig. 7A). The enrichment of bile acid metabolism, xenobiotic metabolism, fatty acid metabolism, heme metabolism and cholesterol homeostasis are encouraging signs of liver-specific organogenesis. Notably, three highly conserved developmental pathways were identified through this analysis – hedgehog, notch and TGFβ. To confirm their functional relevance, we treated organoids with small molecule inhibitors of hedgehog (Cyclopamine – CYC, 0.2µM), notch (DAPT, 10µM) and TGFβR-1 (RepSox, 12.5µM) and characterized the resultant effects on organoid formation. Morphological observations were also correlated with RT-qPCR evaluation of direct transcriptional targets for each signalling pathway (Fig. 7B). Cells managed to established Phase I morphology (where cells lined up the surface of ICC) regardless of the treatment. However, cells treated with RepSox and CYC were unable to form typical organoid structures (Phase II), whilst DAPT treatment appeared to have little effect (Fig. 7C and D). RepSox treated cells arrested in Phase I of organogenesis resembling the observations seen with adult hepatocyte and liver carcinoma cells (**Supplementary Fig. 3)**. CYC treated cells on the other hand, instead of transitioning into typical organoid structures, formed much smaller clusters that were less uniform in size and with a rougher surface. To ensure cytotoxicity was not the causal effect, the selected concentration of each inhibitor used was validated to have had minimal to no significant impact on cell viability (Fig. 7E). Furthermore, the dramatic change in organogenesis observed corresponded to a direct pathway effect (Fig. 7B), reduced hepatic gene expression profile (Fig. 7F), and reduced albumin production rate (Fig. 7G).

**Fig. 7.**
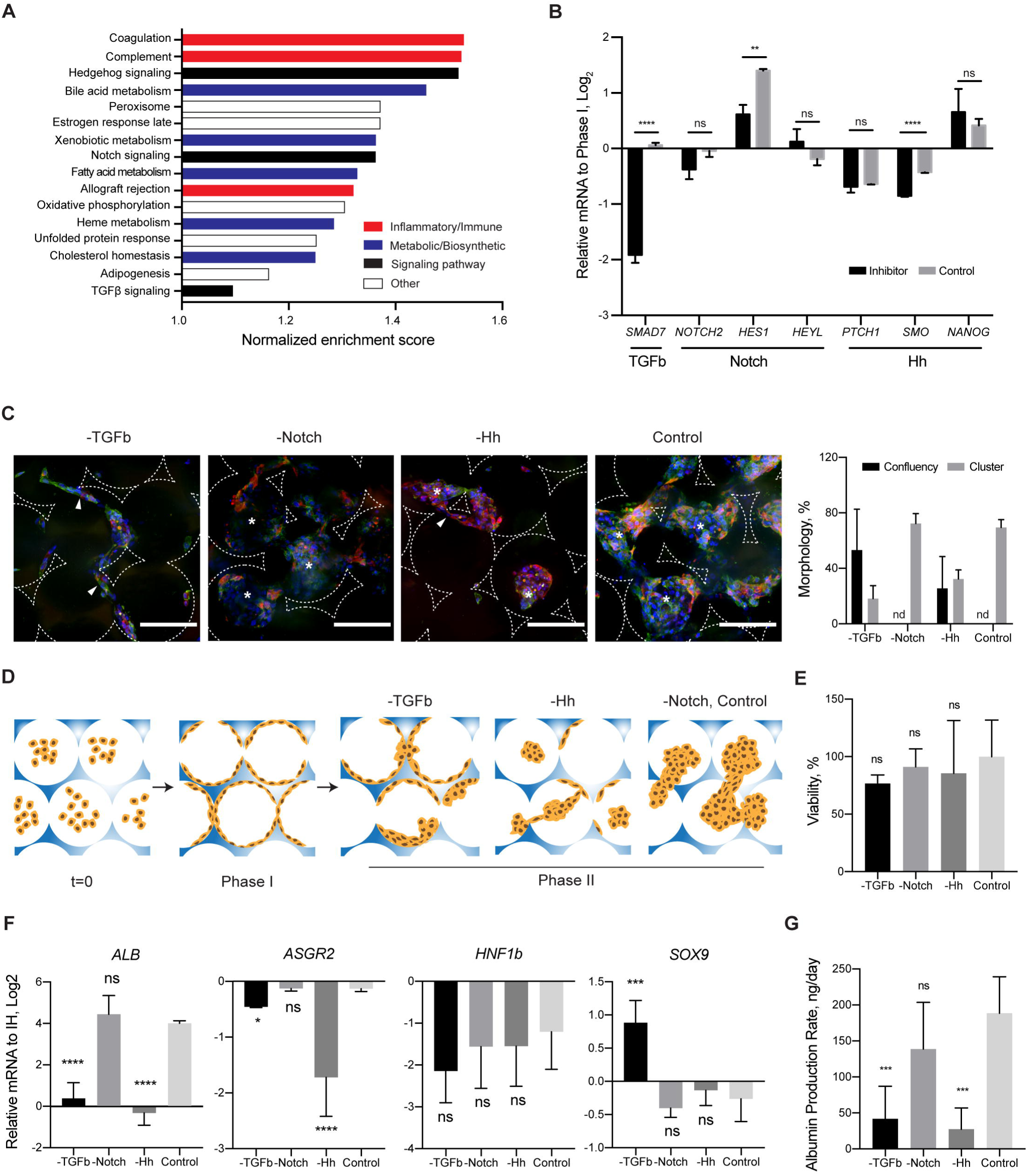
Mechanisms of organoid formation. (**A**) Bar chart detailing gene sets uniquely enriched in IH-ICC over 2D. (**B**) Gene expression (by RT-PCR) in IH-ICC of selected pathway transcriptional targets (TGFb, Notch, Hedgehog) following addition of respective inhibitors (RepSox, DAPT, CYC) (N=4). (**C**) Confocal micrograms of IH in Phase II (CTNNB, green; CK18, red) demonstrating effects of inhibitors on IH-ICC morphogenesis. Morphological quantification of observations provided on right. (**D**) Schematic illustration summarizing effects of inhibitors. (**E**) Cell viability in IH-ICC as a consequence of adding each inhibitor as determined by cell activity (N=8). (**F**) Gene expression (by RT-PCR) of selected hepatic and biliary markers in IH-ICC following addition of each inhibitor above (N=4). (**G**) Effect of inhibitors on IH-ICC hepatic function by albumin production rate (N=4). Mean±sd, ^*^p< 0.05, ^**^p< 0.005, ^***^p < 0.0005, ^****^p<0.0001, ns nonsignificant.

## 4. Discussion

Human organs exist in an incredible variety of shapes and sizes. Understanding the processes through which these varied forms arise during development, are maintained in homeostasis or become perturbed during disease represent some of the most fundamental questions facing us today in biology. iPSC-derived cells provide an excellent model with which to study such questions through ex vivo formation of mini organs known as organoids. iPSC-derived organoids are also highly appealing because they themselves could be used for patient specific disease modeling and therapy. Here we report a new approach to the generation of iPSC-derived hepatic organoids, using state of the art bio-engineering technology that by-passes the need for Matrigel and supporting cell co-culture. Utility of this platform was demonstrated by our findings in organoids of advanced hepatic function and dependence on key developmental signaling pathways (Hedgehog and TGFβ).

Ex vivo liver organogenesis is particularly challenging due to the daunting complexity of the organ’s structure and function. A pivotal breakthrough in this field was made several years ago by Takebe and colleagues when they generated hepatic organoids by co-culturing iPSC-derived hepatic endoderm, HUVECs and MSCs in Matrigel [1]. Immediate downstream translational applications are potentially limited however by dependence upon non FDA-compliant materials such as mouse-sarcoma-derived Matrigel and the difficulty of scaling up production when using multiple cell types (MSCs & HUVECs) [6]. A more recent and equally exciting organoid generation system, developed by Huch & Clevers [2], is similarly limited by the need for Matrigel but also by its need for primary tissue. In this study, we therefore sought to engineer liver organoids free from Matrigel, MSCs, and HUVECs, using a 3D synthetic hydrogel scaffold and hiPSC-derived hepatic progenitors alone. To that end we developed a 3D scaffold called ICC (inverse colloidal crystal) resembling the precise architectural microenvironment into which the liver diverticulum engages during liver bud formation [15]. Due to the ICC’s unique bottom-up fabrication approach, we were able to custom engineer ‘cell-matrix’ interactions through coating the inner lining of the ICC pores with select ECM proteins [15] as well as custom engineer ‘cell-cell’ interactions through manipulating pore size [8]. These engineering designs cumulatively sought to recapitulate the extracellular niche sensed by hepatic progenitors during bud formation. We observed that the initial bud morphology, which we describe in this study as Phase I organogenesis / budding, could only be achieved when two critical factors were met. First, presence of a suitable ECM coating to facilitate cell attachment which otherwise fails due to the bioinert property of the PEG from which the ICC is manufactured (Fig. 2A-D). Second, seeding into a suitably sized pore, which otherwise forces cells to clump rather than attach to the scaffold’s architecture if pore sizes are too small (Fig. 2E-H). Importantly, cell condensation (as opposed to cell-ECM engagement) was associated with down-regulation in hepatic gene expression, lower albumin production rates and failure to progress to Phase II in which cells lining the ICC self-organize into interconnected clusters to create a homogenously repeating organoid tissue structure. This series of morphogenic movements (Phase I → Phase II) was unique to iPSC (IH) and primary (fetal) hepatic progenitors and did not occur with terminally differentiated adult cells or with cancer cell lines (**Supplementary Fig. 3**) [21, 22]. We therefore hypothesize our 3D scaffolds leverage the intrinsic ability of liver stem / progenitor cells to form organized structures in a manner which conserves the interaction observed between progenitors and their surrounding niche during organogenesis / budding [23].

As a consequence of this two-phase organogenesis / budding, the ICC engineered organoids produce structures with graded anatomical distribution of cellular maturity resembling that seen in vivo. Such hepatic zonation as it is more commonly known is previously well documented. In healthy adult liver for example, the functions of hepatocytes exhibit a strict spatial separation regulated by the oxygen gradient along the axis from portal triads to central veins [24]. Zone I periportal hepatocytes have the highest oxygen uptake and are specialized in oxidative liver functions, gluconeogenesis and urea synthesis. Conversely, the perivenous Zone III hepatocytes specialize in glycolysis, lipogenesis and CYP450 drug detoxification. From the perspective of liver differentiation, the maturational lineage is thought to start from Zone I and end in Zone III where the most mature cells are found [25]. The 3D structures seen in our ICC scaffold therefore exemplify a simplistic model of in vivo liver zonation, in which progenitor cells are found at the periphery and more mature cells at the core (Fig. 3A and B). This configuration is in direct contrast to findings seen with spheroid / cell condensation culture systems where cells at the periphery are more viable and proliferative than cells at the core due to hypoxia and DNA damage [26–29]. That difference underscores the success of the engineering feat achieved here. The fact that liver specific drug metabolism and disease modelling become upregulated in parallel with these morphogenic changes along with the capacity of the organoids to integrate, neo-vascularise and survive in vivo emphasizes the physiological relevance of the tissue created here (Fig. 4 and 5). How such morphogenesis and zonation is programmed has been a long-standing question in biology. Our initial interrogation suggests both the TGFβ and hedgehog signalling pathways to be pivotal in this regard. Hedgehog signalling is particularly interesting in the context of liver zonation because it has been reported to orchestrate the position of hepatocytes along an oxygen gradient [24]. Our treatment of progenitors with a hedgehog signalling pathway inhibitor, Cyclopamine, generated not only smaller but dysmorphic organoids (Fig. 6C and D) and was accompanied by downregulation of liver function (Fig. 6F and G). These data suggest Hedgehog may be involved in much more than simple cell positioning therefore.

To our knowledge, this is the first attempt to produce liver organoids using just the combination of iPSC-derived hepatic progenitors and a synthetic hydrogel. The unique modular features of the ICC scaffold will allow study of the complex, combinatorial influences of physical and chemical signals during liver organogenesis in a physiologically relevant, dissectible 3D microenvironment. In addition, the scalable structure and clinically compliant material (PEG) and cell lines [11] used, opens up the possibility of future human therapy. These results highlight the enormous potential of bioengineered organoids for discovery and translational science.

## Data availability statement

The raw/processed data required to reproduce these findings cannot be shared at this time due to technical or time limitations.

## Acknowledgements

We would like to thank Prof Fiona Watt, Simon Broad, Anming Xiong, and Maria Teresa Catanese for reagents and support.

**Author Contribution**
All authors contributed extensively to the work presented in this paper. S.S.N., C.W.F, N.J.C., H.N., J.S.G. and S.T.R. developed the study concept and design. S.S.N., K.S., J.M.S., M. P. S., S.J.I.B., M.H.L. and D.Y.N. acquired data. S.S.N., K.S., J.M.S., J.S.G. and S.T.R. analyzed and interpreted the data. S.S.N., J.S.G. and S.T.R. drafted the manuscript and revised the manuscript for important intellectual content. J.S.G. and S.T.R. supervised the study.

## Abbreviations

iPSC: induced pluripotent stem cell
IH: iPSC derived hepatic progenitors
FDA: food and drug administration
TGFβ: transforming growth factor beta
3D: three-dimensional
ICC: inverted colloidal crystal
PEG: polyethylene glycol
ECM: extracellular matrix
Col: collagen
Fn: fibronectin
Ln: laminin
CK19: keratin 19
ECAM: epithelial cell adhesion molecule
AFP: alpha-fetoprotein
ASGR1: asialoglycoprotein receptor 1
ALB: albumin
HNF4a: hepatocyte nuclear factor 4-alpha
RT-qPCR: reverse transcription-quantitative polymerase chain reaction
2D: two-dimensional
RNA-seq: RNA-sequence
DE: iPSC derived definitive endoderm
PCA: principle component analysis
MRP2: multidrug resistance protein 1
BSEP: bile-salt efflux pump
ZO-1: tight junction protein 1
CLF: cholyl-lysyl-flourescence
TGZ: Troglitazone
HCV: hepatitis C virus
CLDN1: claudin 1
OCLN: occludin
SCARB1: scavenger receptor class B type 1
LDLR: low-density lipoprotein receptor
HSGP: heparin sulfate glycoprotein
H&E: hematoxylin and eosin
CYC: cyclopamine
HUVEC: human umbilical endothelial cell
MSC: mesenchymal stem cell

